# Viral mimicry and memory deficits upon microglial deletion of ATRX

**DOI:** 10.1101/2024.05.07.592875

**Authors:** S. Shafiq, A. Ghahramani, K. Mansour, M. Pena-Ortiz, J.K. Sunstrum, Y. Jiang, M.E Rowland, W. Inoue, N.G. Bérubé

## Abstract

The importance of chromatin-mediated processes in neurodevelopmental and intellectual disability disorders is well recognised. However, how chromatin dysregulation in glial cells impacts cognitive abilities is less well understood. Here, we demonstrate that targeted loss of the ATRX chromatin remodeler in microglia alters chromatin accessibility profiles, leading to the de-repression of endogenous retroelements, triggering viral mimicry. Functionally, we find that ATRX microglial deficiency alters the electrophysiological properties of hippocampal neurons and causes deficits in object recognition and spatial memory. Overall, these findings demonstrate that ATRX is required in microglia to preserve chromatin structure and maintain microglial homeostasis. Disruption of these functions elicit neuroinflammation and cognitive deficits and potentially contribute to the pathology of human neurological disorders caused by *ATRX* mutations.

## Introduction

Microglia are the immune cells of the central nervous system and play critical roles in brain development, homeostasis and disease. They modulate synaptic circuit remodelling, synaptic plasticity, and neuronal activity, which in turn can impact learning and memory (1-3). Microglia are dynamically regulated and can exist in resting/homeostatic or activated/responding states depending on endogenous or exogenous stimuli (4). Whereas resting microglia contribute to nervous system homeostasis and neuronal plasticity through the secretion of neurotrophic factors, microglial states of activation can induce neuronal dysfunction by producing neurotoxic factors and proinflammatory molecules (5, 6). Microglial activation has been observed in neurodegenerative diseases such as Alzheimer’s, Parkinson’s, and Huntington’s diseases and amyotrophic lateral sclerosis, as well as neurodevelopmental disorders including Down syndrome, autism spectrum disorder, and Rett syndrome(7-12). This strongly suggests that active neuroinflammation may account for compromised neuronal survival, synaptic dysfunction and cognitive deficits observed in these pathologies. Indeed, inhibiting the activated state of microglia has been shown to restore neuronal survival and cognitive deficits (10, 13, 14), highlighting the importance of microglia homeostasis in cognitive functions.

ATRX is a Snf2-type chromatin remodeler with crucial functions in the central nervous system (15-17). In humans, *ATRX* mutations cause syndromic and non-syndromic intellectual disability (ID). In its syndromic form (ATR-X syndrome, OMIM #301040), patients also display seizures, microcephaly, hypomyelination and severe developmental delays (18). ID mutations are largely located in two hotspots: the DNMT3a/b and DNMT3L (ADD) globular domain and the Snf2-like enzymatic domain (19, 20). One of the major protein interactors of ATRX is DAXX, which is a histone chaperone for the variant histone H3.3. Together, ATRX and DAXX incorporate H3.3 at repetitive regions of the genome, including telomeres, pericentric repeats, rDNA repeats and endogenous retroviral elements (21-24).

We previously reported that targeted deletion of *Atrx* in postnatal neurons of the mouse forebrain causes long-term spatial and associative memory deficits, in parallel with hippocampal structural alterations and impaired hippocampal synaptic transmission (17, 25). In the present study, we show that *Atrx* deletion in microglia of the mouse central nervous system impacts chromatin accessibility and gene expression profiles in these cells. Our results show that in microglia, ATRX limits DNA damage and the expression of retroelements. Upon loss of ATRX, microglia are activated and trigger the DNA and RNA sensing pathways, leading to an interferon response and cytokine release. These changes are associated with electrophysiological abnormalities in hippocampal CA1 neurons, memory deficits and reduced anxiety. These findings identify a critical role for ATRX and chromatin structure in microglia and the non-cell autonomous deleterious effects on cognitive processes.

## Results

### Evidence of a microglial activated state is observed upon loss of ATRX

We targeted *Atrx* deletion in microglia by mating tamoxifen-inducible *Cx3cr1*^ERT2^ mice (26) and *Atrx*^LoxP^ mice (16), resulting in male progeny lacking ATRX in microglia (ATRX miKO mice). The nuclear membrane Sun1GFP reporter (IMSR_JAX:021039) was also introduced to track the fate of Cre-expressing microglia (27). Cre expression was induced by daily intraperitoneal injections of tamoxifen in 45-day-old mice for five consecutive days (P45-P49) (Figure 1A). There was no difference in weight between control and ATRX miKO mice at 2- and 3-months of age, nor was there a difference in *Cre* and *Sun1GFP* expression by quantitative RT-PCR, suggesting equivalent level of recombination in mice of both genotypes (Figure S1A,B). The efficiency of Cre recombination in microglia was evaluated by quantifying the number of cells expressing Sun1GFP and the microglial marker Ionized calcium binding adaptor molecule 1 (Iba1), revealing over 95% overlap in these markers in control and ATRX miKO mice across multiple brain regions (Figure S1C). Assessment of ATRX protein expression by immunofluorescence staining of brain sections shows that more than 90% of Sun1GFP^+^ cells lack ATRX expression across various brain regions in ATRX miKO mice (Figure 1B,C).

**Figure 1:**
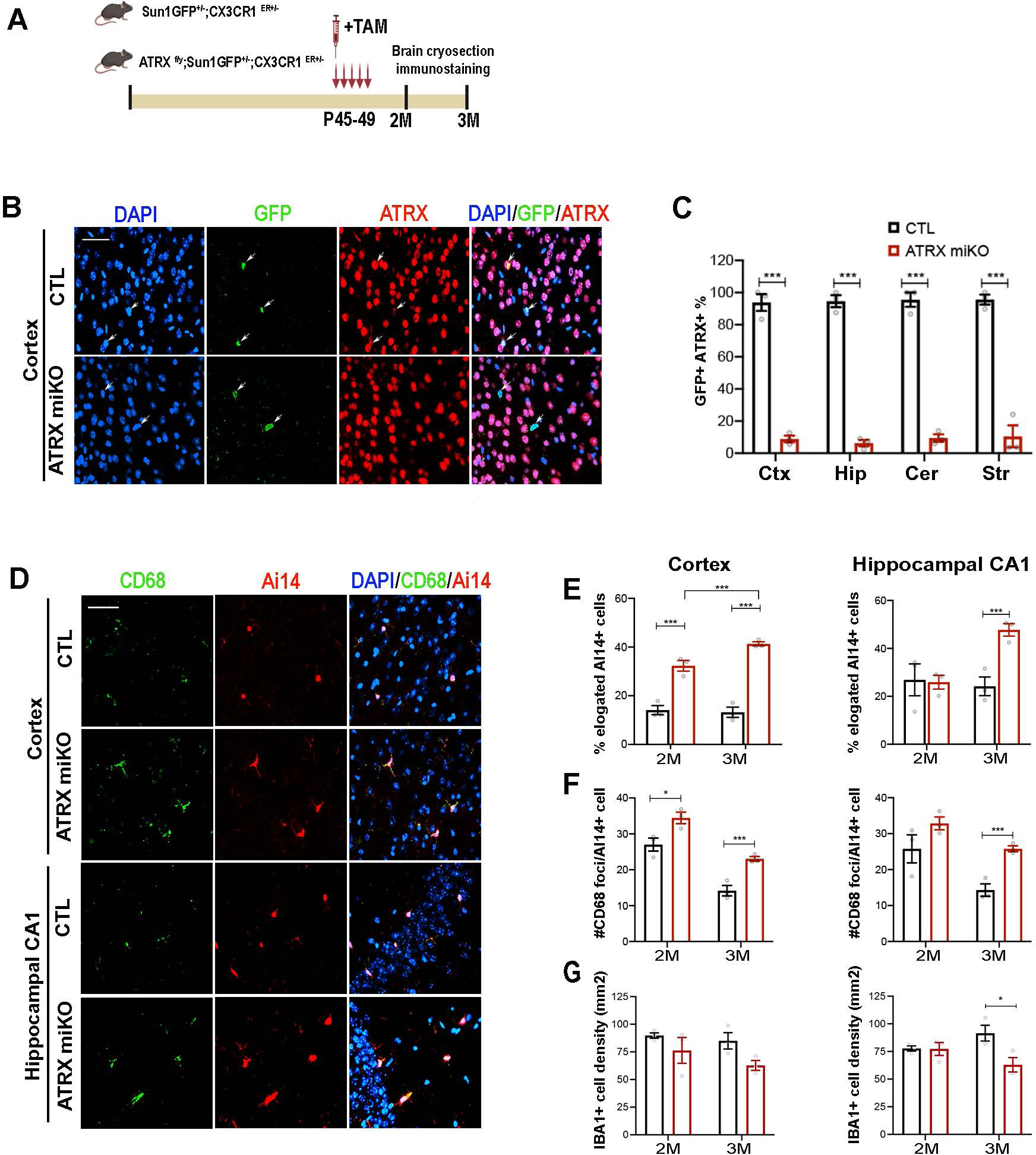
Altered morphology and increased CD68 staining in ATRX-null microglia. (A) Schematic representation of the experimental approach. (B) lmmunofluorescence images of anti-GFP (green) and anti-ATRX (red) counterstained with DAPI (blue) showing absence of ATRX in Sun1GFP+ microglia nuclei in 2 month-old ATRX miKO mice. Scale bar, 50 μm. (C) Quantification of ATRX knockout efficiency in different brain regions (n=3 each genotype, cortex p=0.0001, hippocampus p=0.00003, cerebellum p=0.0001, striatum p=0.0003). (D) lmmunofluorescence staining of CD68 (green) and Ai14+ hippocampal CA1 and cortical microglia in 3 month-old mice. Scale bar, 50 μm. (E) Quantification of microglia with elongated soma in cortex and hippocampal CA1 region of 2 and 3-month-old mice (n=3 each genotype, 2 month CA1 elongated p=0.895, CA1 round p=0.416; 3 month CA1 elongated p=0.007, CA1 round p=0.001; 2 month cortex elongated p=0.003, round p=0.002; 3 month cortex elongated p=0.0002, round p=0.0001). (F) Number of CD68 foci per Ai14+ microglia in cortex and CA1 of 2 and 3 month-old mice (n=3 each genotype, 2 month CA1 p=0.036, 3 month CA1 p=0.005; 2 month cortex p=0.174, 3 month cortex p=0.004). (G) IBA1+ microglia density in cortex and CA1 (n=3 each genotype, cortex 2 month p=0.100, 3 month p=0.00001; CA1 2 month p=0.974, 3 month p=0.0001). Statistical analysis, Student’s T test. Ctx: cortex, Hip: hippocampus, Cer: Cerebellum, Str: striatum.

We next replaced the Sun1GFP with the tdTomato-Ai14 Cre-sensitive allele to allow morphological visualization of control and ATRX-null microglia in brain sections. Upon microscopic examination of tdTomato fluorescence(28), we noticed that microglia lacking ATRX appear morphologically altered compared to control microglia (Figure 1D). We categorized microglia soma structure as round, intermediate or elongated (Figure S1D), and quantified each category in the cortex and hippocampus. This revealed that ATRX miKO mice have an increased proportion of microglia with elongated soma at 2 months, with a larger effect seen at 3 months in the cortex (Figure 1E, S1E,F). Microglia nuclei are larger and appear rod-shaped in ATRX-null compared to control microglia (Figure S1G,H). Hippocampal microglia (CA1, CA2, CA3 and DG regions) also had elongated soma and nuclei, but this only became evident at 3 months of age (Figure 1E, S1 I-K).

We considered that the change in morphology of ATRX-null microglia might reflect an activated state (4), which we evaluated by staining brain cryosections with Cluster of differentiation 68 (CD68)(29). The number of CD68 foci in Iba1+ cells is significantly increased in the cortex of 2- and 3-month-old ATRX miKO mice (Figure 1F). As seen with the change in morphology, increased CD68 staining is only observed at 3 months in hippocampal regions of ATRX miKO mice (Figure 1F, S1L-N). Microglia density is largely maintained, except for a decrease detected at in the 3-month old hippocampus (Figure 1G, S1O,P). Overall, these results indicate that loss of ATRX in microglia alters their morphology and increases CD68 levels, suggestive of an activated state, and that these effects appear earlier in the cortex compared to the hippocampus.

### Transcriptome analysis reveals changes in cell proliferation, genome integrity and the innate immune response

We next evaluated how deleting ATRX in microglia disrupts the transcriptome. Sun1GFP^+^ microglia nuclei were obtained from the cortex and hippocampus of control and ATRX miKO mice using fluorescence-activated nuclei sorting (Figure 2A) (27, 30). RNA-seq of bulk microglia population identified 6168 differentially expressed genes (DEGs) between control and ATRX-null microglia, 3265 of which were upregulated, and 2903 downregulated (Adj p<0.05, Table S1). Gene Set Enrichment Analysis (GSEA) of the upregulated DEGs identified biological process categories mainly related to cell proliferation (sister chromatid segregation, DNA replication, nuclear division, mitotic cell cycle), DNA damage repair (DNA metabolic processes, DNA repair, cellular response to DNA damage stimulus), and to the immune response (defence response) (Figure 2B, Table S2). For downregulated DEGs, the top 10 enriched biological processes are largely linked to synaptic transmission, neuronal morphogenesis, cell adhesion and memory (Figure 2B). GSEA enrichment plots demonstrate that upregulated genes involved in the cell cycle (i.e. *Ccnb1, Ccnd3 E2F2, E2F3, E2F7, E2F8, Cdc25c, Mcm10*), DNA repair (i.e. *Rad51, Brc1a, Fanca, Gen1, DNA2, Blm, DNA2*) and the immune response (i.e. *Cd69, Cd72, Axl, Mx1, Mx2, Tnf, Stat2, Cxcl10*), were significantly over-represented in ATRX-null compared to control microglia (Figure 2C). To confirm these results, we performed immunofluorescence staining for γH2AX, a marker of double stranded breaks and Ki67, a marker of cell proliferation. The results confirm that ATRX-null microglia contain more double-stranded breaks (Figure 2D,E). There was also a notable increase in proliferative Ki67+ ATRX-null microglia in the cortex and hippocampus at 2 months that largely (but not completely) resolved by 3 months of age (Figure 2F,G). We then performed deconvolution of the RNA-seq data to estimate the relative proportions of microglia subpopulations in control and ATRX-null microglia based on published single-cell data(31). Of the nine microglia subclusters identified, the predicted proportion of “activated response” and “interferon response” microglia subclusters are significantly increased in ATRX miKO mice compared to control mice, seemingly at the expense of homeostatic microglia, again suggesting that ATRX deletion causes an immune response in microglia (Figure 2H).

**Figure 2:**
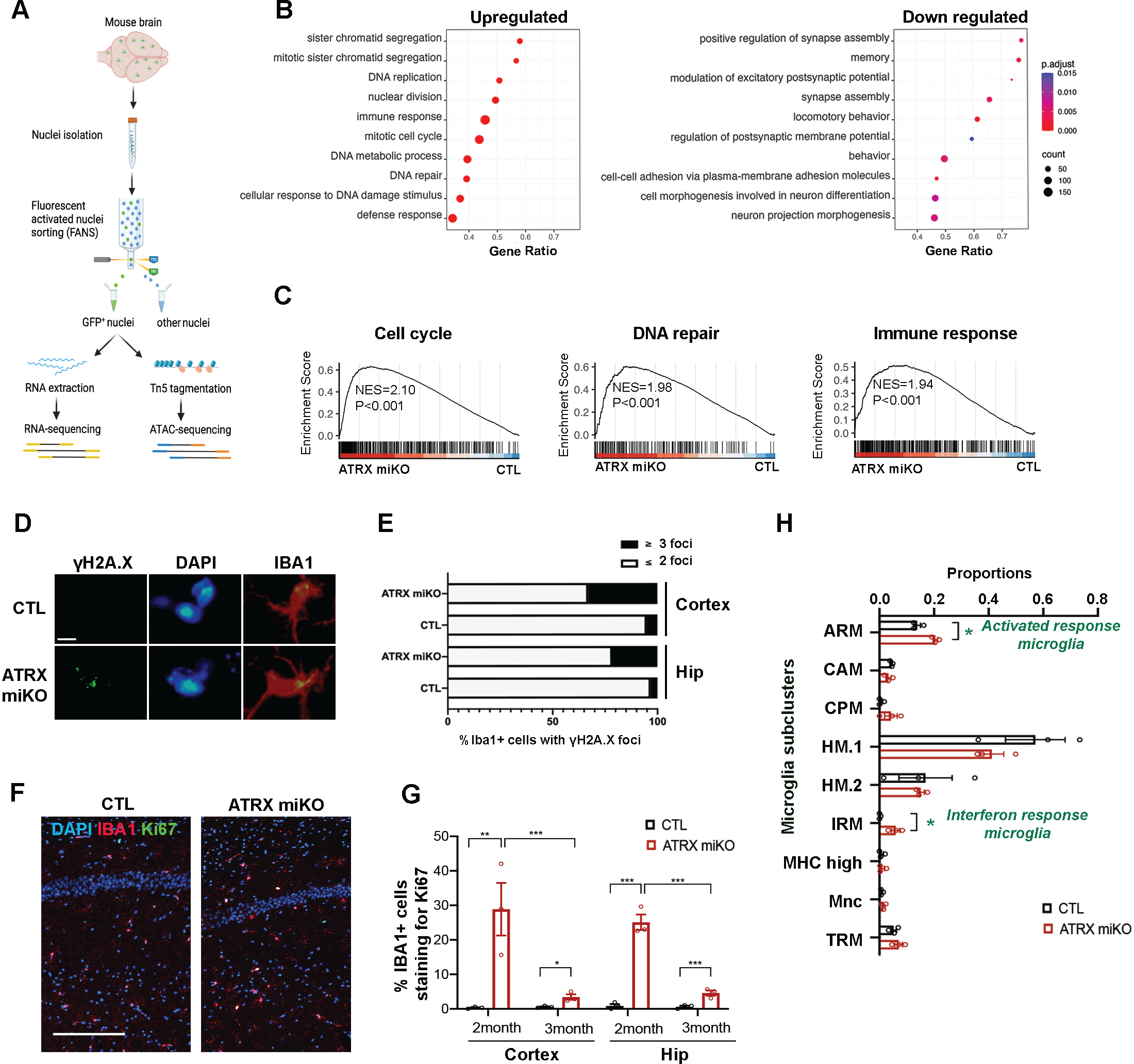
Ablation of ATRX in microglia leads to increased proliferation, DNA damage and immune activation. (A) Schematic illustration of fluorescence-activated microglia nuclei sorting from the mouse cortex followed by RNA-sequencing. (B) The top 10 functionally enriched pathways for upregulated and downregulated genes. (C) Gene set enrichment analysis reveals enrichment of genes linked to cell cycle, DNA repair and immune activation in ATRX deficient microglia. NES, normalized enrichment score. (D) Immunofluorescence staining of IBA1 and γ-H2AX in the cortex of control and ATRX miKO mice. Scale bar, 10mM. (E) Quantification of γ-H2AX foci per microglia in the cortex and hippocampus (n=3 each genotype, cortex p=0.003, hippocampus p=0.018, Student’s T-test). (F) Immunofluorescence staining of IBA1 and Ki67 in the hippocampus of control and ATRXmiKO mice at 2 months of age. Scale bar, 200mM. (G) Quantification of Ki67-positive microglia in the cortex and hippocampus reveals increased proliferation of ATRX-null microglia (n=3 each genotype, Ki67 2 month cortex p=0.020, 3 month cortex p=0.025; 2 month hippocampus p=0.0005, 3 month hippocampus p=0.009, Student’s T-test). (H) Deconvolution of bulk microglia nuclei RNA-seq into single-cell identify sub-clusters. ARM: activated response microglia; CAM: CNS-associated macrophages; CPM: cycling and proliferating microglia; HM.1: homeostatic microglia cluster 1; HM.2: homeostatic microglia cluster 2; IRM: interferon-response microglia; MHC.high: high MHC-expressing microglia; Mnc: monocytes; TRM: transitioning microglia. *p<0.05, Student’s T-test. Error bars represent SEM. The single cell data used for deconvolution was downloaded from GEO data base (GSE142267; Sierksma et. al., EMBO Mol Med 2020, 12, (3), e10606).

### Increased chromatin accessibility in ATRX-null microglia

To investigate whether the changes in transcription upon loss of microglial *Atrx* are associated with altered chromatin accessibility, Sun1GFP^+^ microglia from the cortex and hippocampus were sorted and subjected to the assay for transposase-accessible chromatin followed by sequencing (ATAC-seq) (Figure 2A) (30). Two different approaches, MACS2-DESeq2 and csaw-EdgeR identified a similar number of DARs with comparable genomic distribution (Figure S2A-B). We proceeded with the list of DARs identified using MACS2 for downstream analysis. A total of 32,936 DARs (Adj p<0.05), located within gene promoters, exons, introns, downstream regions and distal intergenic regions were identified (Table S3, Figure S2B), the majority (94%) displaying increased accessibility in ATRX deficient microglia compared to controls (Figure 3A). Accessibility heatmaps and violin plots of DARs emphasize a marked increase in chromatin accessibility at genes, especially at the TSS (Figure 3B,C).

**Figure 3:**
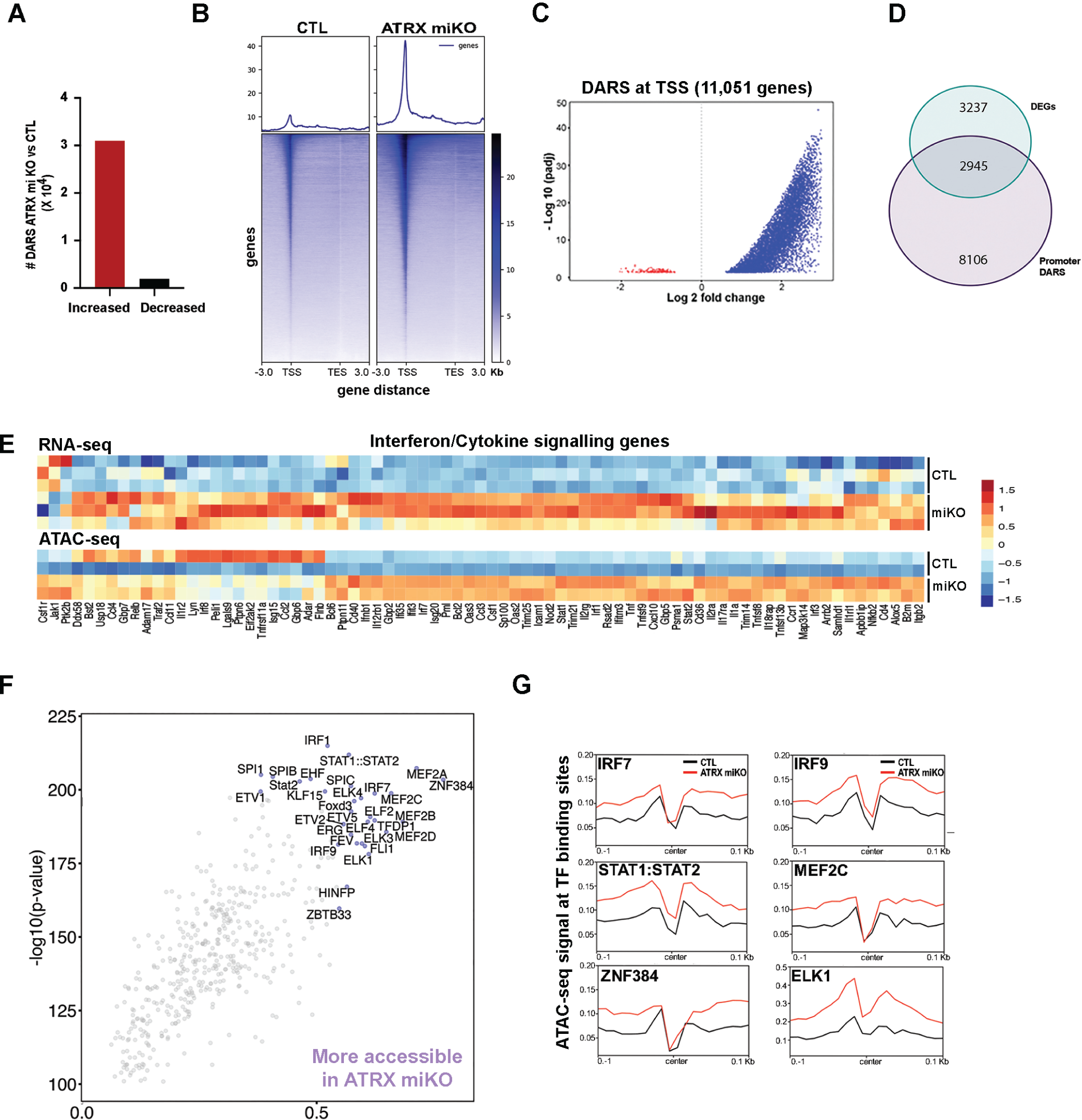
Chromatin accessibility is increased at interferon response/cytokine genes and at interferon signalling TF binding motifs. (A) More genomic regions exhibit increased (red) vs decreased (black) chromatin accessibility in ATRX-null compared to control sorted microglia. (B) Heatmap of chromatin accessibility differences between control and ATRX-null microglia highlights general increased accessibility at promoters and gene bodies in ATRX-null microglia. (C) Violin plot of all genes where DARS are localized at the TSS. (D) Venn diagram showing the extent of overlap between DEGS identified by RNA-seq and DARs at gene promoters. (E) Heatmap of RNA-seq and ATAC-seq data highlights correlation between gene expression and chromatin accessibility changes at interferon and cytokine signalling genes. (F) TF binding motif enrichment analysis from increased DARs. Top 2% are labelled. (G) Genome-wide enrichment of interferon-related TF binding motifs in DARs identified by ATAC-seq. Stronger dips in the center indicate higher confidence in binding and the height of the nearest summit to the center indicates chromatin accessibility. DAR: differentially accessible region, DEG: differentially expressed gene, TSS: transcription start site, TES: transcription end site, CTL: control.

Intersection of the ATAC-seq and transcriptomic data shows that 2945 of the total 6182 DEGs ( ∼47 %) have a DAR at their TSS (Figure 3D), suggesting that altered chromatin accessibility is often coupled with transcriptional changes. We also find that Increased chromatin accessibility is associated with increased expression of genes related to the immune response, cell cycle and DNA repair (Figure 3E, S2C). Within the immune response category, DEGs mainly belong to the innate immune response, the interferon signaling pathway, and the cytokine signaling pathway, and a large majority of DEGs are significantly upregulated and their chromatin more accessible in ATRX-null microglia (Figure 3E).

We next characterized the transcription factor (TF) binding site profiles represented in the ATAC-seq data. We scanned all the annotated TF motifs and scored their differential binding using TOBIAS (32). We identified 30 TFs with increased differential binding scores (top 2%) between control and ATRX miKO (Figure 3F and Table S5). The accessibility of binding sites for TFs linked to interferon signaling, including IRF1, STAT1, STAT2, IRF7, IRF9, ELK1, MEF2C and ZNF384 was increased ATRX miKO (Figure 3G), confirming a broad activation of an immune response in ATRX-null microglia.

### Deletion of ATRX in microglia activates endogenous retroelements

Failure to suppress retroelements in the genome is known to trigger an immune response (33). Given ATRX’s role in heterochromatin formation and previous reports of ATRX binding to retrotransposons in mouse embryonic stem cells (22, 34), we investigated whether activation of the immune response in ATRX-null microglia might be due to the de-repression of retroelements. The analysis of the ATAC-seq data identified 11,155 DARs (Adj p<0.05) in repetitive sequences between control and ATRX-null microglia, with >99% exhibiting increased accessibility (Figure 4A, Table S5). These mainly belong to GC/C-rich repeats, simple repeats as well as long terminal repeats (LTR), especially ERVK and ERVL as well as non-LTR transposable elements (LINE and SINE) (Figure 4B, Figure S2D). From the RNA-seq data, we identified 20,189 differentially expressed retroelements (Adj p<0.05) between control and ATRX miKO (Table S6). Consistent with ATAC-seq, the majority of these were de-repressed in ATRX-null microglia (71%), and mainly belong to ERVK and ERVL types of LTR, as well as LINE and SINE types (Figure 4C,D). This analysis clearly demonstrates that loss of ATRX leads to a marked de-repression of LTR and non-LTR transposable elements in microglia.

**Figure 4:**
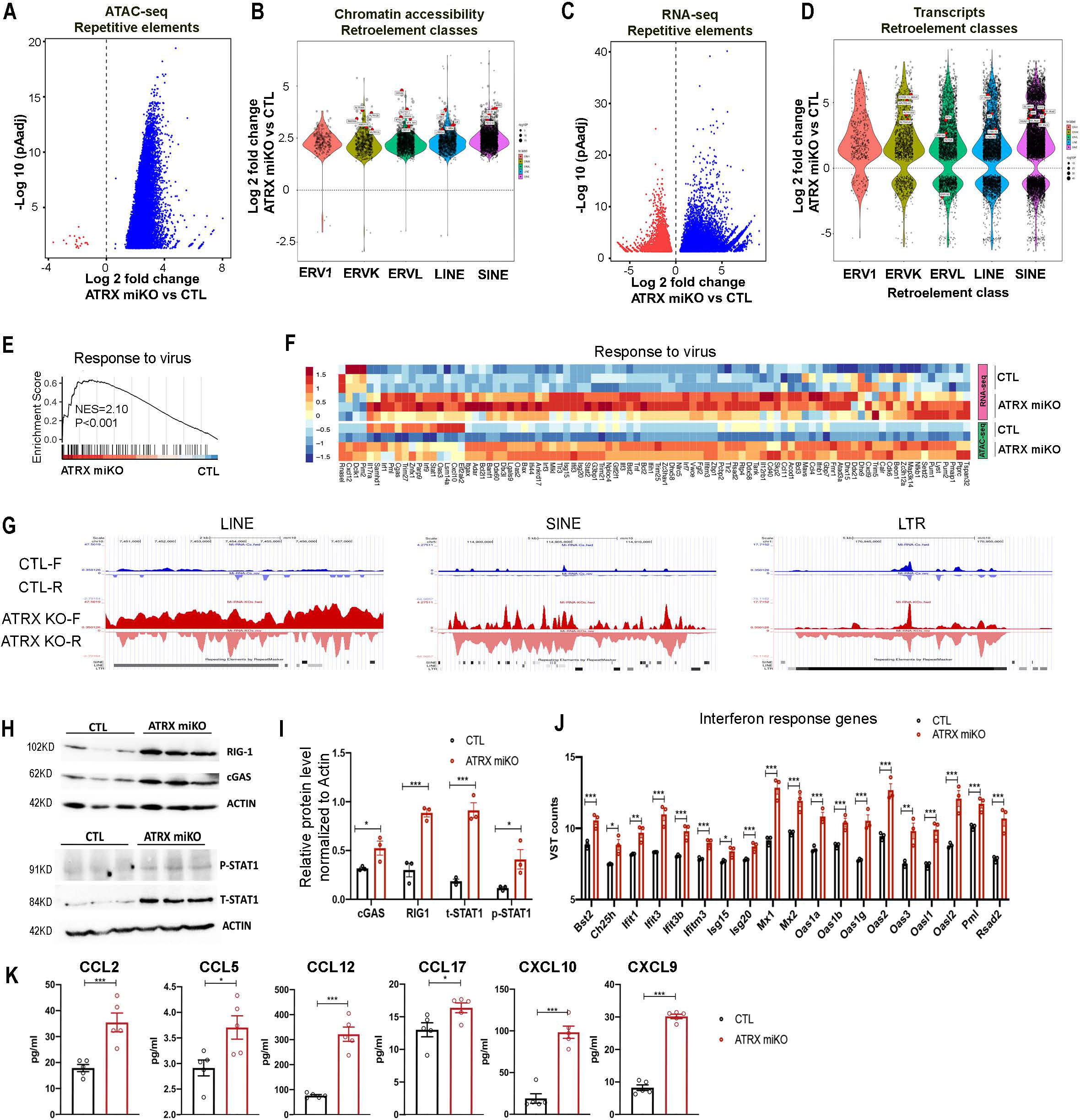
Deletion of Atrx in microglia de-represses endogenous retroelements and triggers the DNA and RNA sensing pathways. (A) Volcano plot of ATAC-seq results for sequence repeats with increased (blue) and decreased (red) chromatin accessibility in ATRX-null microglia. (B) Differential chromatin accessibility between control and ATRX-null microglia for each class of retroelements. (C) Volcano plot showing retroelement loci with increased (blue) and decreased (red) transcript levels in ATRX-null compared to control microglia. (D) Differential expression for each class of retroelements between ATRX miKO and control sorted microglia nuclei. (E) GSEA reveals that the “response to virus” pathway is significantly enriched in ATRX-null microglia. NES, normalized enrichment score. (F) Heatmap representing transcript levels and chromatin accessibility for the “response to virus” pathway. (G) UCSC genome browser tracks showing bi-directional transcription of retroelements belonging to LINE, SINE and LTR in sorted ATRX miKO microglia nuclei. F and R represent forward and reverse strands, respectively. (H) Western blots of DNA/ RNA sensing pathway components and (I) graph of relative protein levels (n= 3 each genotype; cGAS p=0.047, RIG-1 p=0.002, pSTAT1 p=0.042, STAT1 p=0.001, Student’s T-test). Error bars represent +/-SEM. (J) Gene expression of interferon stimulated genes (ISGs). Variance stabilizing transformation (VST) counts were taken from RNA-seq data (n= 3 each genotype; Bst2 p=0.007, Ch25h p=0.022, Ifit1 p=0.012, Ifit3 p=0.003, Ifit3b p=0.006, Ifitm3 p=0.004, Isg15 p=0.037, Isg20 p=0.009, Mx1 p=0.006, Mx2 p=0.006, Oas1a p=0.001, Oas1b p=0.007, Oas1g p=0.002, Oas2 p=0.002, Oas3 p=0.015, Oasl1 p=0.003, Oasl2 p=0.004, Pml p=0.007, Rsad2 p=0.006, Student’s T-test). Error bars represent +/-SEM. (K) Cytokine/chemokine levels (n= 5 each genotype; CCL2 p=0.001, CCL5 p=0.020, CCL12 p=0.000, CCL17 p=0.032, CXCL10 p=0.000, CXCL9 p=0.000, Student’s T-test).

### Loss of ATRX engages a viral mimicry and interferon response

The expression of genes related to a viral response was significantly enriched in ATRX-null microglia (Figure 4E) and correlated with increased chromatin accessibility (Figure 4F). Both de-repression of bidirectional retroelements and DNA damage have been shown to trigger a viral mimicry response through sensing of cytosolic RNA or DNA (35-38). Bidirectional transcription of retroelements could result in the formation of double-stranded RNA (dsRNA) (39, 40), which trigger the interferon response (41, 42). Strand-specific analysis of the RNA-seq data indeed shows an increase in both the sense and antisense retroelement transcripts upon *Atrx* deletion in microglia (Figure 4G). Combined with the observed increase in DNA damage in ATRX-null microglia (Figure 2D,E), we considered that one or both of these might engage the viral mimicry pathway. Indeed, multiple genes belonging to the DNA/RNA sensing pathways are overexpressed in ATRX-null microglia (from the RNA-seq data, data not shown), including Cyclic GMP-AMP synthase (cGAS) and Retinoic acid-inducible gene I (RIG-1). cGAS senses double-stranded DNA (dsDNA) and produces cyclic guanosine monophosphate–adenosine monophosphate (cGAMP), which then binds Stimulator of interferon response CGAMP interactor 1 (STING1) (43). Western blot analysis shows that the cGAS protein is elevated in the ATRX miKO cortex, suggesting engagement of the DNA sensing pathway (Figure 4H,I). RIG-1 senses double-stranded RNA (dsRNA) and interacts with the Mitochondrial antiviral-signalling protein (MAVS) (43). Western blot analysis shows increased RIG-1 in the ATRX miKO cortex, indicating the potential activation of the RNA sensing pathway (Figure 4H,I).

To confirm that one or both of these sensing pathways is engaged, we next examined the state of common downstream effectors. The cGAS or RIG-1 signaling cascades result in the phosphorylation and heterodimerization of the Signal transducer and activator of transcription 1 and 2 (STAT1, STAT2) (44). We found that total STAT1 (t-STAT1) and phospho-STAT1 (p-STAT1) protein levels are increased in ATRX miKO cortical tissue compared to controls (Figure 4H,I). Downstream of STAT activation, the expression of interferon stimulated genes (ISGs) was increased in ATRX-null microglia (Figure 4J).

Recognition of nucleic acids by pattern-recognition receptors (PRRs), such as RIG-1 and cGAS, is essential for triggering an antiviral immune response by inducing the production of proinflammatory cytokines (45). A cytokine/chemokine array analysis of cortical protein extracts revealed an increase of several cytokines/chemokines in ATRX miKO mice, including CC motif ligand 2 (CCL2), CC motif ligand 5 (CCL5), CC motif ligand 12 (CCL12), CC motif ligand 17 (CCL17), C-X-C motif chemokine ligand 9 (CXCL9) and C-X-C motif chemokine ligand 10 (CXCL10)(Figure 4K and Table S7). Together, these results support the idea that loss of ATRX in microglia causes activation of the DNA and RNA-sensing pathways and the interferon response.

### Loss of microglial ATRX impacts electrophysiological states of hippocampal CA1 neurons

Microglia sense neural activity and in turn modulate neuronal activity (46). Therefore, we assessed the impact of microglial ATRX loss on the neuronal membrane properties (i.e. intrinsic properties) as well as excitatory synaptic transmission in hippocampal CA1 pyramidal neurons. We first studied the subthreshold properties by artificially injecting a small-amplitude current in whole-cell current clamp recordings (Figure 5A). We found that CA1 neurons from ATRX miKO mice exhibit a lower membrane resistance (input resistance) compared to control mice (Figure 5B). A decrease of the input resistance can be due to an increase in the cell size (surface membrane area (47)). However, the membrane capacitance (Cm), a proxy for the surface membrane area, was similar between the ATRX miKO and control (Figure 5C), suggesting that CA1 neurons lacking ATRX have increased opening in membrane channels with little, if any, change in cell size. In response to the steps of depolarizing current injections, we observed a regular-spiking pattern as reported elsewhere for hippocampal pyramidal neurons(48) in both control and ATRX miKO mice (Figure 5D). The overall current-spike frequency relationship is similar between control and ATRX miKO mice (Figure 5E). Other firing properties, including latency and inter-spike interval, are also not different between control and ATRX miKO mice (Figure 5F and data not shown). The examination of the action potential shapes elicited by the minimal current injection (rheobase) revealed that CA1 neurons from ATRX miKO mice exhibit an increased action potential amplitude compared to control mice (Figure 5G,H). The rheobase current and threshold were not different between control and ATRX miKO neurons (Figure 5I,J). We also studied spontaneous excitatory postsynaptic current (sEPSC) in voltage-clamp recordings (Figure 5K). The frequency and inter-event distribution (Figure 5L and not shown) were not different between control and ATRX miKO mice. However, ATRX miKO mice showed a significant leftward shift in the sEPSC amplitude distribution (Figure 5M). The average sEPSC amplitude change did not reach a statistically significant decrease (Figure 5N), suggesting that the synaptic change may occur in subpopulations of synapses and their amplitude changes may be masked when expressed as population average. Together, this data demonstrates that microglial ATRX is required to maintain normal electrophysiological properties of hippocampal CA1 pyramidal neurons.

**Figure 5:**
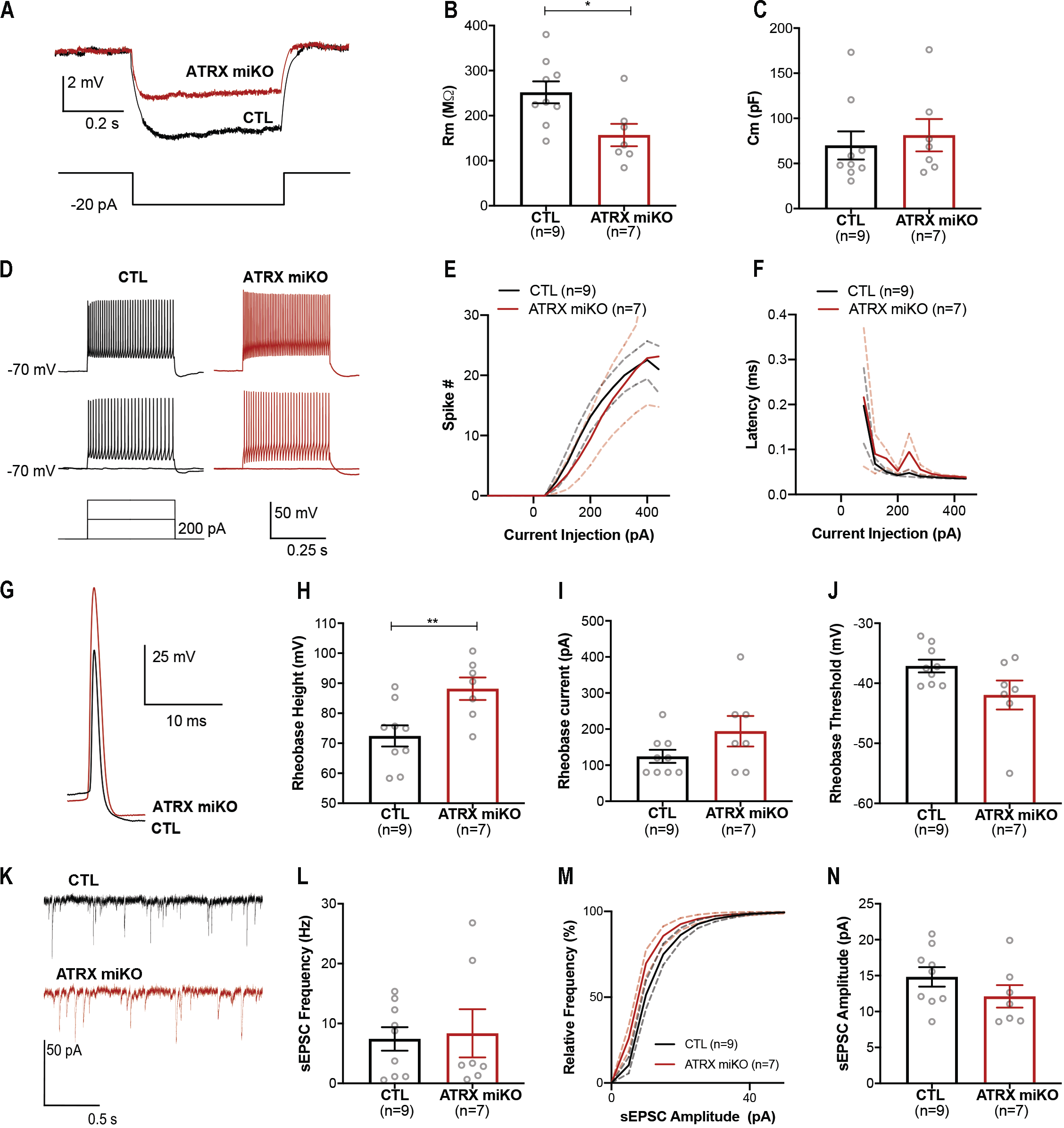
Altered electrophysiological properties of CA1 hippocampal pyramidal neurons in ATRX miKO mice. (A) Representative current clamp traces of -20pA step. The step protocol is shown at the bottom. (B-C) Summary of membrane resistance (Student’s T-test, p=0.017) and capacitance (Student’s T-test, p=0.3266) calculated from the -20pA step in A. (D) Representative current clamp traces of control (left, black) and ATRX miKO (right, red) firing during 400pA (top), 200pA (middle) and 0pA current steps from a holding potential of -70 mV. The step protocol is shown in the bottom left. (E) Average spike number (CTL n=9, ATRX miKO n=7, two way ANOVA, F (1.000, 8.000)=0.07102, p=0.796). The solid line represents the current injection curve, and the dotted lines represent SEM. (F) Latency to the first spike in the rheobase sweep plotted by the current step (two way ANOVA, F(1, 8)=2.608, p=0.145). The dotted lines represent SEM. (G) Representative traces of rheobase spike in control and ATRX miKO neurons. (H-J) Summary of action potential amplitude (Student’s T-test, p=0.009), rheobase current (Student’s T-test, p=0.1212) and threshold (Student’s T-test, p=0.068). (K) Representative voltage clamp traces showing sEPSCs with 100uM picrotoxin to block GABA-sIPSCs. -70mV holding potential. (L) Summary of sEPSC frequency (Student’s T-test, sEPSC frequency p=0.8277). (M-N) Cumulative distribution of sEPSC amplitude (Kolmogorov-Smirnov test, p < 0.0001) and summary of sEPSC amplitude (Student’s T-test, sEPSC amplitude p=0.2100). Error bars represent SEM.

### Loss of ATRX in microglia leads to memory deficits and anxiolytic effects in mice

We next assessed whether the molecular and cellular defects observed in ATRX-null microglia impact cognitive abilities. In the open field test, ATRX miKO and control mice travel an equivalent distance (Figure 6A), indicating that general locomotor activity is not affected. The total time spent in the centre also did not differ between genotypes (Figure 6B).

**Figure 6:**
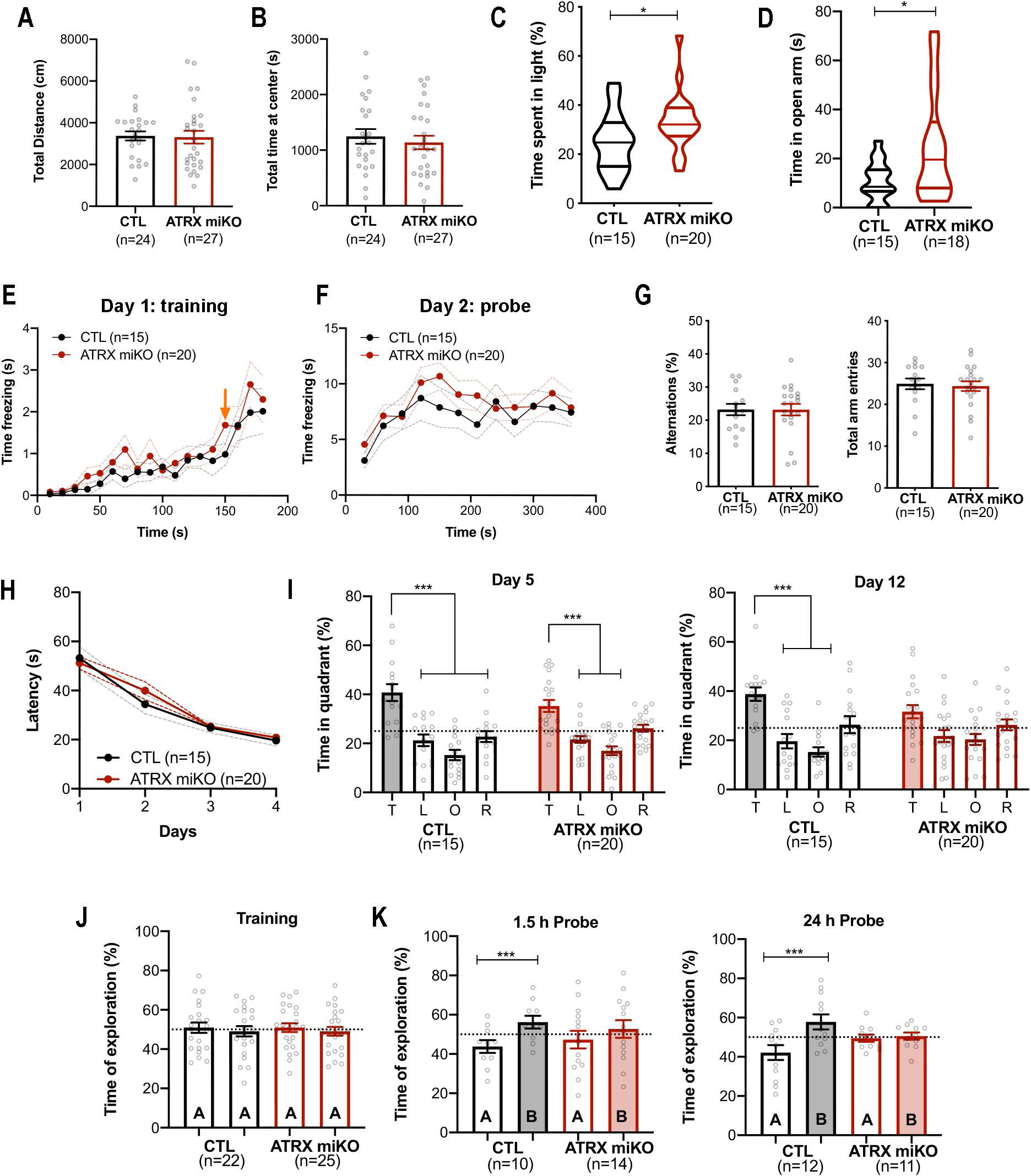
Spatial and object recognition memory impairments in ATRX miKO mice. (A) Total distance travelled in the open field test (CTL n=24, ATRX miKO n=27, p=0.884, Student’s T-test).(B) Total distance travelled in the center of the open field (CTL n=24, ATRX miKO n=27, p=0.553, Student’s T-test). (C) Percent time spent in the light chamber during the light-dark box test (CTL n=15, ATRX miKO n=20, p=0.039, Student’s T-test). (D) Percent time spent in the open arm of the elevated plus maze test (CTL n=15, ATRX miKO n=18, p=0.032, Student’s T-test). (E) Time freezing over 180s during the training phase of the contextual fear conditioning task. Arrow indicates the shock administered at 150s. (F) Time freezing over 360s during the contextual fear conditioning probe test (CTL=15, ATRX miKO=20, F(1, 33)=0.6785, p=0.416, two way ANOVA). (G) Percent sponteneous alternations and number of intersections over 5 min in the Y-maze task (CTL n=15, ATRX miKO n=20, percent alternations p=0.982, total arm entries p=0.761, Student’s T-test). (H) Latency to reach the platform over 4 days of training (4 trials/day) in the Morris water maze paradigm (CTL n=15, ATRX miKO n=20, F(1, 132)=0.4290, p=0.5136, two way ANOVA). (I) Percent time spent in the target (T), left (L), right (R) and opposite (O) quadrants after the removal of the platform on probe day 5 (Ftarget (2.036, 67.20)=28.15, Fgenotype(1, 33)=0.04870, ***p= <0.001, two way ANOVA), and at probe day 12 (Ftarget(2.370, 78.22)=13.05, Fgenotype(1, 33)=1.960, ***p= <0.001, two way ANOVA). The dotted lines indicate random chance. (J) Percent of time spent exploring object A during the 10 min training session in the novel object paradigm (CTL p=0.721, ATRX miKO, p=0.673, Student’s T-test). (K) Percent of time spent exploring object A and a new object B at 1.5h (CTL n=10, p=0.0151, ATRX miKO n=14, p=0.400, Student’s T-test) and 24h after training (CTL n=12, p=0.008, ATRX miKO n=11, p=0.686, Student’s T-test). The dotted lines indicate random chance. Error bars represent ± SEM.

Several anxiety tests were conducted, revealing noteworthy differences between the ATRX miKO and control groups. In the light-dark paradigm, ATRX miKO mice spend significantly more time in the lit area compared to control mice, suggesting reduced anxiety levels (Figure 6C). This finding is further supported in the elevated plus maze test during which ATRX miKO mice spend significantly more time in the open arm than control mice (Figure 6D). However, there is no significant difference between groups in the number of entries into different arms of the elevated plus maze (Figure S3A).

In the contextual fear conditioning test, ATRX miKO mice spend the same amount of time freezing as control mice (Figure 6E,F) and in the Y-maze, the percent spontaneous alternations and total arm entries were not different between genotypes (Figure 6G), indicating that contextual fear memory and working memory processes are intact in ATRX miKO mice.

In the Morris water maze task, ATRX miKO and control mice had equivalent latency to find the platform as well as speed and distance travelled over the four days of training (Figure 6H, S3B,C). Short-term and long-term spatial memory were tested on day 5 and day 12, respectively. On day 5, both control and ATRX miKO mice spend significantly more time in the target quadrant than the left, opposite, or right quadrant, indicating intact short-term spatial memory in ATRX miKO mice. However, on day 12, ATRX miKO mice did not spend significantly more time in the target quadrant, indicative of hippocampal-dependent long-term spatial memory deficits (Figure 6I). Finally, mice were evaluated in the novel object recognition memory test. During the habituation period, both control and ATRX miKO mice equally explored the two identical objects (Figure 6J). However, when memory was tested at 1.5h and 24h later, whereas ATRX miKO did not show a preference for the novel object (Figure 6K), indicating an impairment in short and long-term recognition memory. In summary, behavioural testing shows that mice lacking *Atrx* in microglia exhibit lower anxiety levels and deficits in spatial and object recognition memory.

## Discussion

ATRX is a chromatin remodeling protein linked to intellectual disability that plays a crucial role in gene regulation and chromatin organization, and in maintaining genomic integrity. In this study, we find that targeting *Atrx* deletion in microglia increases chromatin accessibility, DNA damage and de-represses retroelements, triggering viral mimicry pathways and an interferon-mediated immune response. Functionally, we demonstrate that microglial-specific ATRX inactivation alters the electrophysiological properties of hippocampal CA1 neurons and leads to memory deficits and anxiolytic effects.

Our findings demonstrate that ATRX-mediated heterochromatin formation is required to fully suppress retroelements in microglia. The ATAC-seq data exposes a marked global de-condensation of chromatin, especially at gene TSS and retrotransposons, concomitant with aberrant transcriptional activation of both ERVs and non-LTR retrotransposons. These retroelements are also de-repressed in a variety of neurodegenerative, neurological and psychiatric disorders (49-52), and ERV activation has been linked to immune response, nucleic acid sensing response, immuno-inflammation, structural changes in hippocampal pyramidal neurons, and cognitive deficits (53, 54). Specific repression of ERVs has been shown to rescue cognitive deficits (54), suggesting that retroelement activation may have contributed to the neuroinflammation and cognitive deficits observed upon *Atrx* deletion in microglia (53, 54). The deletion of ATRX in oligodendrocytes or astrocytes does not cause such an extensive de-repression of retroelements as observed in microglia (data not shown), suggesting that epigenetic regulation of retroelements in microglia might be different and more susceptible to ATRX loss. Several layers of suppression have been described to keep retroelements in check, including DNA methylation, H3K9me3 and H3K27me3 (55, 56). ATRX can promote both H3K27me3 and H3K9me3 based on reports of its interaction with EZH2 and SETDB1, respectively (57). Future studies could address whether microglia have relatively low DNA methylation at retroelements, which would provide an explanation for enhanced vulnerability of ATRX-null microglia.

We provide evidence that ATRX is required cell-autonomously in microglia to maintain genomic stability. The latter drives several human diseases, such as cancer, neurodegeneration, and early aging (58-60). We previously discovered that loss of ATRX in mouse neuroprogenitor cells leads to replication stress, DNA damage and mitotic defects (61, 62). It is therefore not that surprising to detect increased DNA damage in ATRX-null microglia, which could be exacerbated by the increased proliferation of activated microglia detected at 2 months of age. DNA damage in ATRX-null microglia could potentially generate cytoplasmic DNA species and trigger the DNA sensing pathway (63),(64). In addition to DNA damage, we detected bi-directional transcription of ERVs as well as non-LTR retrotransposons that can generate immunogenic small dsRNAs and R-loops (39, 40),(65). These can be recognized by PRRs, such as cGAS or RIG-1, and trigger interferon-mediated immune responses (41, 42). Given that we observe upregulation of both RIG-1, cGAS, and the downstream outcomes (proinflammatory cytokines and chemokines production, interferon-stimulated gene upregulation), we propose that the effects observed in the ATRX miKO mice might stem from the combined activation of the DNA and RNA sensing pathways.

Our findings also highlight the non-cell autonomous impact of ATRX-null microglia on neurons. The loss of ATRX expression in microglia led to a significant decrease in the input resistance in CA1 pyramidal neurons. The lower input resistance generally indicates hypo-excitability of the neurons because, by Ohm’s law (*V*=*I* x *R*), a given positive ion influx (*I*) by excitatory synaptic transmission results in a smaller postsynaptic depolarization (*V*) with lower input resistance (*R*). On the other hand, microglial ATRX deficiency also led to an increase in the action potential amplitude. While the classic view is that action potential transmits signals in all- or-none fashion (48), a growing body of evidence shows that changes in the action potential amplitude can fine-tune the Ca^2+^ dynamics at the axon terminals and the ensuing neurotransmitter release (66-68). Thus, our data suggests that microglial ATRX deficiency may change signal transmission from CA1 pyramidal neurons to their downstream targets. Mechanistically, an increase in action potential amplitude can be explained by an increase in the voltage-gated Na+ channels that drives depolarization and/or a decrease in voltage-gated K+ channels that counteract the depolarization. Future studies on the precise biophysical changes in CA1 pyramidal neurons will be required to clarify the mechanisms by which microglia influence neuronal function.

It is not yet clear which of the identified alterations in microglia is responsible for cognitive deficits displayed by ATRX miKO mice. When activated, microglia can secrete proinflammatory cytokines and chemokines that influence synaptic plasticity and cognitive functions. Excessive amounts of dsRNA can activate microglia and release cytokines/chemokines, such as CCL2, CLL5, CXCL9 and CXCL10 (69-74), as we observe in ATRX-null microglia. Overproduction of these cytokines and chemokines has previously been demonstrated to modulate neuronal activity and cause cognitive deficits(73, 75-80). Moreover, dsRNA generated from the bi-directional transcription of retroelements may be linked to the anxiolytic effects observed upon *Atrx* deletion in microglia (73, 75-81).

Our study also highlights the cell type-specific contributions of ATRX in supporting cognitive functions. Microglia-specific deletion of *Atrx* causes short- and long-term recognition memory deficits not previously observed from a deletion in postnatal excitatory neurons (17). Similarly, we observed fear conditioning memory deficits upon *Atrx* deletion in postnatal neurons (17), but not in the ATRX miKO model. These differences demonstrate that a neuro-centric view of cognition underestimates the complexity of the nervous system and limits our ability to progress towards efficient therapies for neurodevelopmental disorders. In that regard, our findings indicate that cognitive deficits in patients harboring *ATRX* mutations could stem in part from microglial activation and neuroinflammation. Several potential approaches could be considered to limit these effects, for example by repressing endogenous retroelements (54), attenuating nuclei acid sensing (53), neutralizing chemokines/cytokines (82), or suppressing microglial immune activation (10, 13, 14).

## Material and methods

### Animal husbandry and genotyping

Mice were exposed to the 12-hour light/12-hour dark cycles and fed regular chow and water ad libitum. ATRX^loxP^ mice were previously described (16), and two reporter lines (Sun1GFP (B6;129-Gt(ROSA)26Sor^tm5(CAG-Sun1/sfGFP)Nat^/J, MGI:5614796, RRID: IMSR_JAX:021039) and Tomato-Ai14 (Ai14) allele (B6.Cg-Gt(ROSA)26Sor^tm14(CAG-tdTomato)Hze^/J, MGI:3809524, RRID:IMSR_JAX:007914)) were bred with the ATRX^loxP^ mice (27, 28). Upon Cre-mediated recombination, Sun1GFP green fluorescent protein will label the nuclear membrane, while Tomato-Ai14 will express tdTomato protein with red fluorescence in the cytoplasm and nucleus. Male progeny with *Atrx* deficiency in microglia upon tamoxifen treatment was produced by mating ATRX^loxP^;Sun1GFP or ATRX^loxP^;Tomato-Ai14 females to males expressing Cre under the inducible promoter of Cx3Cr1^ER^ (B6.129P2(Cg)-Cx3cr1^tm2.1(cre/ERT2)Litt^/WganJ, MGI:5617710, RRID:ISMR JAX:021160) (26). Genomic DNA was extracted from an ear notch and genotyped as described previously (30). The genotyping primers are listed in Table S8. Mouse weight was checked with weighing balance at 2 months and 3 months of age. Behavioral tests were performed using male mice of 3 to 6 months of age, starting from less demanding to more demanding ones (open field tests, light-dark box, elevated plus maze, Y-maze, novel object recognition, fear conditioning and Morris water maze). All behavioral tests were performed on at least 15 animals per genotype between 9:00 AM and 4:00 PM. ARRIVE guidelines were followed: mouse groups were randomized, experimenters were blind to the genotypes, and software-based analysis was used to score mouse performance in all the tasks. The Animal Care and Use Committee of the University of Western Ontario approved all animal procedures in compliance with the Animals for Research Act guidelines of the province of Ontario, Canada (AUPs 2021-049 and 2021-063).

### Tamoxifen administration

Tamoxifen (10mg; Cat#T5648, Sigma) was dissolved in 100µl 95% ethanol at 65°C for 10 minutes and then diluted with 900µl corn oil (Cat# C8267, Sigma). 45 day-old adolescent male mice were injected intraperitoneally daily with 2mg tamoxifen for 5 consecutive days.

### Immunofluorescence

2-month or 3-month-old mice were transcardially perfused and fixed as described previously (83). Fixed brains were then sectioned, coronally or sagitally, at 8μm thickness (Leica CM 3050S) on Superfrost slides (Thermo Fisher Cat# 22-037-246) and stored at -80°C with a desiccant (VWR, 61161-319). For the immunofluorescence, slides were rehydrated in 1xPBS for 5 min followed by washing with wash buffer (1xPBS + 0.3 % TritonX-100 (Millipore Sigma Cat# T8787)). Antigen retrieval was performed (except for CD68 staining) by incubating slides in 10 mM sodium citrate at 95°C for 10 min. Cooled sections were washed in 1x PBS and then blocked with 5 % goat serum (Millipore Sigma Cat# G9023) or donkey serum (Millipore Sigma Cat# D9663) in wash buffer and incubated with primary antibody overnight at 4°C. After washing 3 times (5 min each) with wash buffer, slides were incubated with secondary antibody for 1 hour at room temperature in the dark. Slides were washed twice for 5 min with washing buffer, and counterstained with 1 μg/mL DAPI (Millipore Sigma Cat# D9542) for 5 min followed by 1x PBS wash. Finally, sections were mounted with Permafluor (Thermo Fisher Cat# TA-006-FM) and imaged with an inverted microscope (DMI 6000b, Leica) equipped with a digital camera (ORCA-ER, Hamamatsu). Volocity (PerkinElmer Demo Version 6.0.1, RRID:SCR_002668) and Adobe Photoshop was used for image processing. All cell counts were performed in a blinded and randomized manner. Primary antibodies used for immunofluorescence were anti-ATRX (Santa Cruz Biotechnology, Cat# sc-15408, RRID:AB_2061023), anti-CD68 (Biorad, Cat#MCA1957, RRID:AB_322219), anti-Ki67 (abcam, Cat#ab15580, RRID:AB_443209), anti-IBA1 (Cedarlane, Cat# 019-19741, RRID:AB_839504), anti-γH2AX (Cell Signaling Technology Cat# 2577, RRID:AB_2118010) and anti-GFP (Thermo Fisher Scientific, Cat# PA1-9533, RRID:AB_1074893). The following secondary antibodies were used: goat anti-rabbit-Alexa Fluor 594 (1:800, Thermo Fisher Scientific, A-11012, RRID:AB_2534079), goat anti-rabbit-Alexa Fluor 488 (1:800, Thermo Fisher Scientific Cat# A-11008, RRID:AB_143165), donkey anti-sheep-Alexa Fluor 594 (1:800 Thermo Fisher Scientific Cat# A-11016, RRID: AB_2534083), goat anti-chicken-Alexa Fluor 488 (1:800, Thermo Fisher Scientific Cat# A-11039, RRID:AB_2534096) and goat anti-rat-Alexa Fluor 488 (1:800, Thermo Fisher Scientific, A-11006, RRID:AB_2534074).

### Analysis of microglia cell morphology

Tomato-Ai14 (Ai14) reporter mice (B6.Cg-Gt(ROSA)26Sor^tm14(CAG-tdTomato)Hze^/J, MGI:3809524, RRID:IMSR_JAX:007914)) were bred with the ATRX^loxP^ mice that expressed tdTomato protein with red fluorescence in the cytoplasm and nucleus in a Cre-dependent manner. Microglia were classified based on the shape of soma, i.e., round, elongated, and medium. For microglia density, all the microglia were labelled with Iba1 immunostaining and the number of Iba1+ microglia were counted and transformed to Iba1+ cells per mm^2^.

### Fluorescence-activated sorting of microglia nuclei

Fluorescence-activated nuclei sorting (FANS) was performed as described previously (30). The cortex and hippocampus of 2-month-old mice were homogenized in 20 mM Tricine KOH, 25 mM MgCl2, 250 mM sucrose, 1 mM DTT, 0.15 mM spermine, 0.5 mM spermidine, 0.1% IGEPAL-630, 1x protease inhibitor cocktail (Millipore Sigma Cat# 11873580001), 1 μL/mL RNase inhibitor (Thermo Fisher Scientific Cat# 10777019)). After dilution with homogenization buffer and filtering through a 40 μm strainer (Fisherbrand Cat#22363547), the samples were layered on the top of 1:1 volume cushion buffer (0.5 mM MgCl2, 0.88 M sucrose, 0.5 mM DTT, 1x protease Inhibitor cocktail (Millipore Sigma Cat# 11873580001), 1 μL/mL RNase inhibitor (Thermo Fisher Scientific Cat# 10777019)). Nuclei were then pelleted at 2800 g for 20 min at 4°C and incubated for 10 min in sorting buffer (4% FBS, 0.15 mM spermine, 0.5 mM spermidine, 1x protease inhibitor cocktail (Millipore Sigma Cat# 11873580001) and 1 μL/mL RNase inhibitor (Thermo Fisher Scientific Cat# 10777019) in 1x PBS). After resuspending and filtering through a 20 μm strainer (PluriSelect Cat#431002060), nuclei were sorted using a Sony SH800 Cell sorter and Sun1GFP+ microglia nuclei were collected into a 1.5ml tube.

### RNA purification and RNA-seq library preparation

RNA extraction followed by RNA-seq library preparation was performed as described previously (30). Briefly, sorted Sun1GFP+ microglia nuclei were collected directly in lysis buffer (supplement with 2% β-mercaptoethanol) from the single cell RNA purification kit (NorgenBiotek Cat#51800). RNA extraction, with on-column DNase treatment, was performed by following the manufacturer’s instructions. 35ng of total RNA was used to deplete rRNA (Ribo-off rRNA Depletion kit (H/R/M), Vazyme Cat#N406), followed by strand-specific RNA-seq library preparation using the VAHTS Universal V8 RNA-seq Library Prep Kit for Illumina (Vazyme Cat#NR605-01).

### RNA-seq analysis

RNA-seq libraries were sequenced at Canada’s Michael Smith Genome Sciences Centre (BC Cancer Research, Vancouver, BC, Canada) using the Illumina HiseqX (Illumina Inc., San Diego, CA), and 60-120 million paired-end reads (150 bp) were obtained for each library. RNA-sequencing analysis was performed as described previously (30, 83). Briefly, Trim galore v0.6.6 with the following parameters (‘–phred33 -length 36 -q 5 -stringency 1 -e 0.1‘) was used to trim the raw data, and HISAT2 version 2.0.4 (84) mapped the paired-end reads against the *Mus musculus* GRCm38.p6 (primary assembly downloaded from Ensembl). SAMtools (85) was then used to sort and convert SAM files. StringTie v.2.1.5 (86) was used to obtain gene and transcript abundance for each sample by providing read alignments and *Mus musculus* GRCm38 genome annotation as input. Tximport R/Bioconductor package was used to import transcript coverage and abundance into R, and the DESeq2 R/Bioconductor package (87) conducted a differential analysis of transcript count data. The independent hypothesis weighting (IHW) method was used to weigh P values and adjust for multiple testing using the procedure of Benjamini Hochberg (BH). Finally, the Lancaster method was used to aggregate transcripts’ p-values. Gene ontology and gene cluster analysis were performed as described previously (83). For Gene Set Enrichment Analysis (GSEA), significant DEGs were ranked based on log2FoldChange. gseGO function from clusterProfiler v4.0.5 (88) was then used with the following parameters (ont = “BP”, OrgDb = org.Mm.eg.db, minGSSize = 10, maxGSSize = 500, eps = 1e-10, pvalueCutoff = 0.05, pAdjustMethod = “BH”) to perform GSEA. For single-cell deconvolution, reference-based decomposition mode from the R toolkit, Bisque v1.0.5 (89), was used to estimate cell composition from our bulk expression data with microglia single-cell data downloaded from GSE142267 (31).

### Assay for transposase-accessible chromatin (ATAC)

ATAC-seq on sorted nuclei was performed as described previously (30). Briefly, sorted Sun1GFP+ microglia nuclei were collected directly in 1x PBS and pelleted at 3000 rpm for 5 min at 4°C. The nuclei were resuspended directly in tagmentation buffer containing Tn5 transposase (Vazyme Cat#TD501) and the reaction mixture was placed at 37°C for 45 min. After tagmentation, samples were purified using the QIAquick PCR Purification Kit (Qiagen Cat#28104) and amplified by PCR according to the TruePrep DNA kit instructions (Vazyme Cat#TD501). Quantitative PCR was performed to determine the optimal number of cycles (1/3 saturation), and the libraries were amplified with no more than 12 PCR cycles.

### ATAC-seq analysis

On average, 100M paired-end reads (150bp) were obtained for each library. Raw reads were trimmed with Trim Galore (v.0.6.6) with the following parameters (‘–phred33 -length 36 -q 5 - stringency 1 -e 0.1‘) and mapped to *Mus musculus* GRCm38.p6 using Bowtie2 v2.4.4 (90) with the default parameters. SAMtools v1.12 was then used to create and sort BAM files from the aligned reads recorded in SAM format. The duplicated reads were then marked with the MarkDuplicates function from Picard (v.2.26.3). The mitochondrial DNA reads, and blacklist regions of the genome were filtered out using Bedtools intersect (v.2.30.0). Two approaches were employed to identify differentially accessible regions (DARs). First, MACS2 v2.2.7.1 (91) (with parameters --nomodel -f BAMPE) was used to call the peaks for each sample. The peaks from all samples were merged into a set of non-redundant open regions with Bioconductor package soGGi v1.24.1. The peaks that were present in at least two samples were kept for downstream analysis. The featurecounts function from Rsubread v2.6.4 (92) was used to count the reads from each sample overlapping the consensus peak set and DESeq2 v1.32.0 was used to perform a differential analysis of accessible regions between ATRX miKO and control samples. Second, R package csaw (v.1.26.0) (93) was used to count the reads in 200bp non-overlapping windows. Background noise was estimated by counting reads in 2000bp bins. We selected 200bp windows that have a signal higher than log2(3) above the background. Windows were then merged if less than 100bp apart but did not extend above 5kb width. EdgeR 3.40.2 (94) was then used to identify the DARs. Because MACS2 and csaw have more than 93 % overlap in calling DARs (Figure S2B), only MACS2 was later used for downstream analysis. DARs (P value < 0.05) in the genome were annotated using ChIPseeker v1.28.3 (95). For TF footprinting, TF motifs were downloaded from the JASPAR CORE database (96). The footprinting analysis was performed using merged BAM files of each condition and a merged set of peaks from all samples. We used TOBIAS v0.12.11 (32) to assess chromatin occupancy by TFs. TOBIAS ATACorrect was used to correct Tn5 insertion bias in input BAM files. Footprinting scores were then calculated using TOBIAS ScoreBigWig. We then used TOBIAS BINDetect to analyze the differential binding of transcription factors between ATRX miKO and control groups.

### Intersection of RNA-seq and ATAC-seq

To generate heatmaps of expression levels, gene-level counts from the Stringtie results were transformed using the variance stabilizing transformation (VST) method in DESeq2. For the heatmap of chromatin accessibility, ATAC-seq signals at promoters of the genes (1000bp upstream and 1000bp downstream of TSS) were counted using the featureCounts function from Rsubread v2.6.4. Z-scores were calculated for expression or accessibility independently. Heatmaps were plotted using the Pheatmap package.

### Analysis of repetitive elements

For the analysis of retroelements, the repeatmasker annotation file for mm10 was downloaded from the UCSC table browser. Reads falling within the repeatmasker annotation were counted for both RNA and ATAC assays using the featureCounts function from Rsubread v2.6.4. Differential expression and accessibility of repeats were determined using DESeq2. For the bi-directional transcription of retroelements, mapped reads from RNA-seq data in BAM format were split into sense and antisense strand BAMs using a custom script. Reads falling within the UCSC repeatmasker annotation were counted for both strands separately using the featurecounts function from Rsubread. Bigwig tracks were generated using Deeptools11 bamCoverage v.3.5.2 with the parameters “-bs 25 –normalize using RPKM” and visualized in the UCSC genome browser.

### Western blot analysis

Total protein was extracted from the cortex and hippocampus of 2 month-old mice using the RIPA buffer (150 mM NaCl, 1% NP-40, 50 mM Tris pH 8.0, 0.5% deoxycholic acid, 0.1% SDS, 0.2 mM PMSF, 0.5 mM NaF, 0.1 mM Na3VO4, 1x protease inhibitor cocktail (Millipore Sigma Cat# 11873580001). Homogenized samples were incubated on ice for 30 min and centrifuged at 12000 rpm for 15 minutes at 4°C. Bradford assay (BioRad Cat# 500-0006) was used to quantify the protein and 100 μg of protein was resolved on 10% SDS-PAGE gel. After transferring proteins to nitrocellulose membrane (BioRad Cat# 1620115), the membrane was blocked with milk-TBST (5% skimmed milk, 1xTBS and 0.1 % Tween-20) and then incubated with anti-STAT1 (Cell Signaling Technology Cat# 9172, RRID:AB_2198300), anti-pSTAT1 (Cell Signaling Technology Cat# 9167, RRID:AB_561284), anti-RIG-1 (Cell Signaling Technology Cat# 3743, RRID:AB_2269233), anti-cGAS (Cell Signaling Technology Cat# 31659, RRID:AB_2799008), anti-Actin (Sigma-Aldrich Cat# A2066, RRID:AB_476693) antibodies at 4°C overnight. After 3 washes with milk TBST, the membrane was incubated with the appropriate secondary antibody (rabbit anti-HRP (1:5000, Jackson ImmunoResearch Cat# 111-036-003, RRID:AB_2337942); mouse anti-HRP (1:5000, Santa Cruz Cat# sc-516102, RRID:AB_2687626)). Finally, the membrane was incubated in an enhanced chemiluminescent solution (Thermo Fisher Cat# 34095) and exposed using the Universal Hood III (BioRad Cat# 731BR00882). Quantification of blots was performed with ImageJ (version 1.53).

### Cytokines/Chemokines Analysis

Total protein was extracted from the cortex and hippocampus of 2 month-old mice using the RIPA buffer and the multiplexing analysis was performed using the Luminex™ 200 system (Luminex, Austin, TX, USA) by Eve Technologies Corp. (Calgary, Alberta). Forty-five markers were simultaneously measured in the samples using Eve Technologies’ Mouse Cytokine 45-Plex Discovery Assay® which consists of two separate kits; one 32-plex and one 13-plex (MilliporeSigma, Burlington, Massachusetts, USA). The assay was performed according to the manufacturer’s protocol. Assay sensitivities of these markers range from 0.3 – 30.6 pg/mL for the 45-plex.

### Slice Preparation for Electrophysiology

Mice were deeply anesthetized using sodium pentobarbital (100mg/kg intraperitoneally) and transcardially perfused with cold (2–4 °C) NMDG-HEPES solution (92mM NMDG, 93mM HCl, 2.5mM KCl, 1.2mM NaH2PO4, 30 mM NaHCO3, 20mM HEPES, 25mM Glucose, 5mM sodium ascorbate, 2mM Thiourea, 3mM sodium pyruvate, 10mM MgCl2, 0.5mM CaCl2 (300–310 mOsm), saturated with 95% O2/5%CO2). Brains were quickly removed and placed in a cold NMDG-HEPES solution for slicing. Coronal sections (350 μm thick) containing the hippocampus were cut using a vibratome (VT-1200, Leica Biosystems). Slices were incubated at 34°C for 15 minutes in NMDG-HEPES solution saturated with 95% O2/5%CO2. Slices were then transferred to artificial cerebrospinal fluid (aCSF) (126mM NaCl, 2.5mM KCl, 26mM NaHCO3, 2.5mM CaCl2, 1.5mM MgCl2, 1.25mM NaH2PO4 and 10mM D-glucose (295–300 mOsm)), saturated with 95% O2/5%CO2 and maintained at room temperature until recording.

### Electrophysiology measurements

Slices were transferred to a recording chamber superfused with aCSF at a flow rate of 1.5–2.0 mL/min and maintained at 27–30 °C. CA1 neurons were visualized using an upright microscope with infrared differential interference contrast optics (BX 51WI, Olympus). Borosilicate glass recording pipettes (BF120-69-15, Sutter Instruments) were pulled in a Flaming/Brown Micropipette Puller (P-1000, Sutter Instruments) with a resistance between 3–5 MΩ. Pipettes were filled with an internal solution (116mM K-gluconate, 8mM KCl, 12mM Na-gluconate, 10mM HEPES, 2mM MgCl2, 4mM K2ATP, 0.3mM Na3GTP and 1mM K2-EGTA (283–289 mOsm, pH 7.2–7.4)). Spike firing was measured in the current clamp from a holding potential of -80mV using a step protocol from -160pA to 460pA in 40pA increments. Glutamatergic sEPSCs were isolated by adding picrotoxin (100 μM) to the aCSF while holding the postsynaptic neuron at -80 mV in a voltage clamp. Access resistance was monitored throughout the recording and cells were discarded if the value exceeded 20 MΩ.

### Data collection and analysis of electrophysiological measurements

Whole cell patch clamp recordings were obtained using a Multiclamp 700B amplifier (Molecular Devices, California, USA), low pas filtered at 1 kHz and digitized at a sampling rate of 20 kHz using Digidata 1440A (Molecular Devices). Data was recorded on a PC using pClamp 10.6 (Molecular Devices) and analysed using MiniAnalysis (Synaptosoft, Georgia, USA) for EPSCs, Clampfit (Molecular Devices) for membrane potential and a custom Python code (adapted from a white paper from Allen Cell Types Database: https://github.com/AllenInstitute/ipfx) for cell firing. Briefly, the slope (dV/dt) was measured by taking the difference in voltage between two time steps and dividing it by the resolution of acquisition. The time of the threshold crossing was detected by finding the time point where dV/dt was ≤5% of the maximum dV/dt of the rising phase. The following criteria were used to detect action potentials during current injection steps: 1) the duration from threshold to the peak is ≤5 ms, 2) amplitude is ≥2mV and absolute peak ≥-30mV, and 3) action potential trough (minimum membrane potential in the interval between the peaks of two consecutive action potentials) is ≤–22 mV. For sEPSC analysis, baseline data was taken at least 5 mins after breaking through into whole-cell mode and a 0.5- or 1-min bin was used for analysis. To achieve an accurate measure of the amplitude, individual sEPSCs were visually screened in MiniAnalysis and events below 5 pA were not included.

### Open field test

The mice were acclimated to the room for 30 minutes prior to testing. Then the mice were placed in an open arena (length 20cm, width 20cm, height 30cm) for 2 hours and locomotor activity was measured in 5 min intervals as previously described (17). Locomotor activity was automatically recorded (AccuScan Instrument), and distance travelled, and time spent in the center were reported.

### Y maze test

The mice were acclimated to the room for 30 minutes prior to testing. The mice were placed in the center of a symmetrical three-armed Y maze as described previously (17) and their activity was recorded for 5 minutes using AnyMaze. The order and number of entries into each arm were reported. Spontaneous alternations were counted when a mouse entered all three arms in a row without visiting a previous arm.

### Light dark box

The mice were acclimated to the room for 30 minutes prior to testing and were then placed in a light-dark box and allowed to explore the arena for 10 minutes. Their activity was automatically recorded (AccuScan Instrument), and the percent time spent in light was reported.

### Elevated plus maze

The mice were acclimated to the room for 30 minutes prior to testing and then placed in the center of the elevated plus maze (Med Associate Inc) and their activity was recorded for 5 min using AnyMaze. Time spent in the open, closed or center of the elevated plus maze was reported. The center of the mouse body was used as an indicator to determine its presence in an open, closed or center.

### Novel object recognition

The mice were habituated for two consecutive days in an empty arena (40cm x 40cm) for 5 minutes as described previously (17). The next day, mice were trained by exposing them to two identical objects (A) for 10 minutes and their activity was recorded using AnyMaze. After 1.5 hours of training, one of the objects was replaced with a novel object (B), and mice were exposed to both old (A) and novel (B) objects to test their short-term recognition memory. Similarly, mice were exposed to objects (A) and (B) 24 hours after training to test their long-term recognition memory. Time spent with objects was recorded when mice sniff or touch the objects but did not lean against and/or climb on the object.

### Morris water maze

The Morris water maze test was conducted as described previously (17), where the mice were placed in a 1.5m diameter pool with 25°C water. Spatial cues were displayed around the pool and the platform was submerged 1cm below the water’s surface. The mice were acclimated to the room for 30 minutes prior to testing and then trained to find the platform in four trials (90 s) a day for 4 consecutive days with a 15 minutes intertrial period. Their activity was recorded with AnyMaze. If the mice did not find the platform within the 90s, they were gently guided onto the platform. Short or long-term spatial memory was tested by removing the platform on day 5 or day 12, respectively. The latency to find the platform over 4 days of training, and the percent time spent in each quadrant of the maze were reported for the probe tests.

### Contextual fear conditioning

The mice were acclimated to the room for 30 minutes prior to testing and then placed in a 20 cm x 10 cm enclosure with a metal grid floor connected to a shock generator. One wall of the enclosure was marked with a stripe pattern and mouse activity was recorded using AnyMaze. For training, mice were placed in the enclosure and allowed to freely explore it. After 150 s, a shock was administered (2 mA, 180 V, 2 s), and the mice were returned to their home cage after 30 seconds. After 24 hours, contextual fear memory was tested by placing the mice in the enclosure for 6 min. Freezing time was reported in 30 s intervals, where freezing was defined as immobility lasting more than 0.5 s.

### Statistical analysis

The Student’s T-test (unpaired, two-tailed) or one-way ANOVA for experiments with one variable or two-way repeated-measures ANOVA with post hoc test for experiments with two variables were performed using GraphPad Prism software (GraphPad Software Inc, California, USA). All results are depicted as mean ± SEM unless indicated otherwise. P values of less than 0.05 were considered to indicate significance. All statistical details are outlined in the figure legends.

## Data availability

The accession number for the RNA/ATAC sequencing data reported in this paper can be accessed at SRA accession number PRJNA787973. Data analysis and graphical representations were performed using R scripts and publicly available packages as denoted in the methods detail section. All scripts are available upon request.

## Acknowledgements

We are grateful to Vania Prado and Marco Prado for the Cx3Cr1^ER^ mice and Samuel Asfaha for access to the Sony SH800 sorter. The behavioural assays were performed at the Robarts Research Institute neurobehavioral core facility and next-generation sequencing was performed at Canada’s Michael Smith Genome Sciences Centre (BC Cancer Research, Vancouver, BC, Canada). This work was supported by the Canadian Institutes of Health Research operating grant to N.G.B. S.S. received a Children’s Health Research Institute Trainee award funded by the Children’s Health Foundation. K.M. received a Pediatrics Summer Studentship and a Pediatrics Graduate Studentship from the Department of Pediatrics at Western University.

## Author contribution

S.S. contributed to conceptualization, investigation, formal analysis, and writing - original draft and review & editing; A.G. contributed to formal analysis and draft review & editing; K.M. and J.S. contributed to investigation, formal analysis, investigation and draft review & editing; M.P., M.R., and Y.J. contributed to investigation and formal analysis, and draft review & editing; W.I. contributed to conceptualization and review & editing. N.G.B contributed to conceptualization, funding acquisition, supervision, and writing original draft and review & editing.

## Supplementary Figure legends

**Figure S1:**
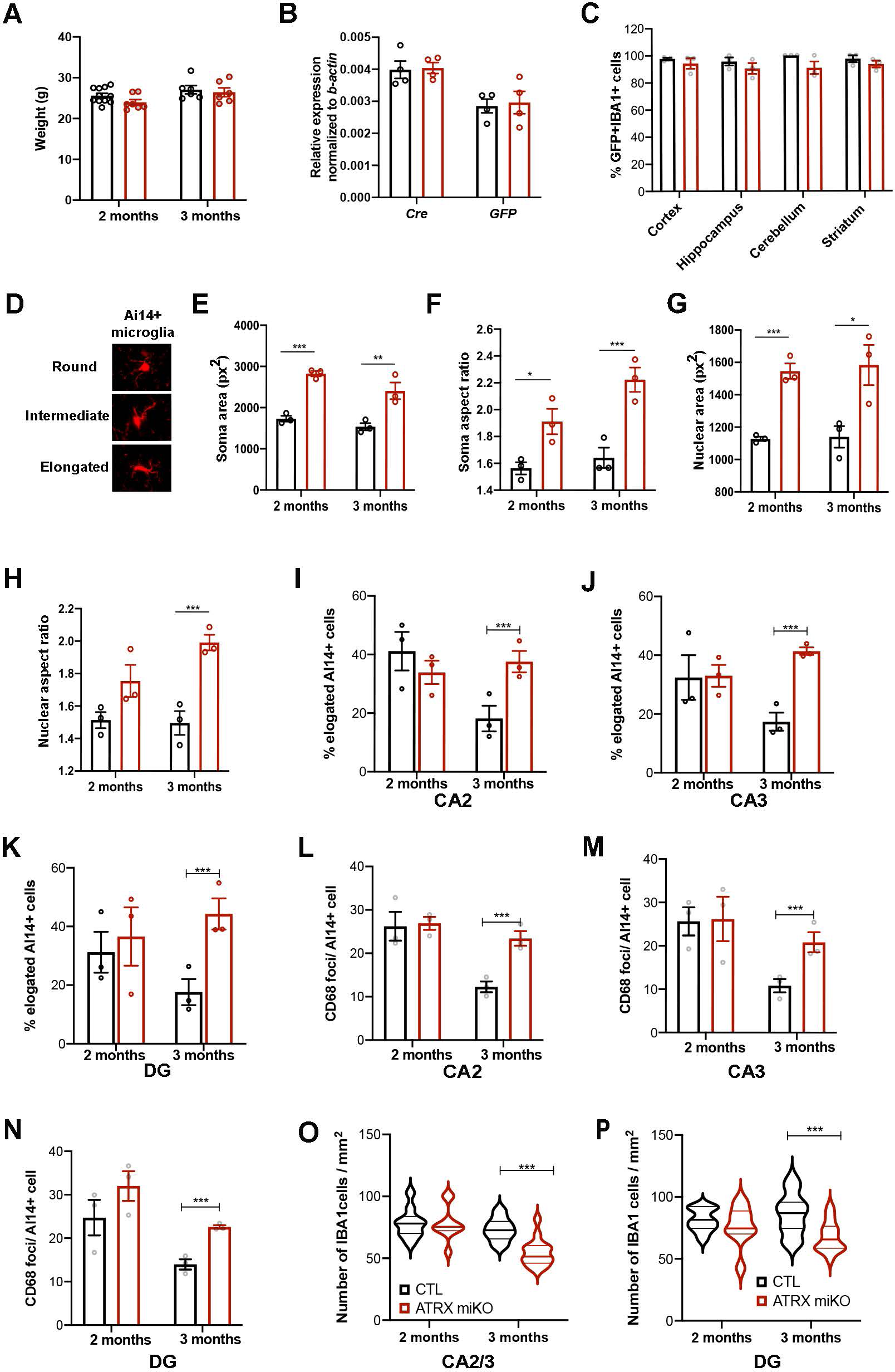
Activation state and morphology of control and ATRX-null microglia. (A) Weight of 2 and 3 month-old control and ATRX miKO mice (2 months, CTL n=11, ATRX miKO n=7, p=0.080; 3 months, CTL n=6, ATRX miKO n=6, p=0.679 Student’s T-test). (B) RT-qPCR of Cre and Sun1GFP transcripts in 2 month-old control and ATRX miKO mice (n=4 each genotype, Sun1GFP p=0.871; Cre p=0.804, Student’s T-test). Results were normalized to beta-actin transcript levels. (C) Quantification of immunofluorescence staining of Sun1GFP and IBA1 reveals >95% Cre expression in control and ATRX miKO mice across different brain regions (n=3 each genotype, cortex p=0.423, hippocampus p=0.351, cerebellum p=0.119, striatum p=0.327). (D) Representative images of AI14+ cells with round, medium and elongated soma. (E-F) Quantification of soma area and aspect ratio in the cortex at 2- and 3-months of age (n=3 each genotype, soma area 2 months p=0.0003, 3 months p=0.016; aspect ratio 2 months p=0.029, 3 months p=0.007, Student’s T-test). (G-H) Quantification of nuclear area and aspect ratio in the cortex at 2- and 3-months of age (n=3 each genotype, nuclear area 2 months p=0.001, 3 months p=0.034; aspect ratio 2 months p=0.0937, 3 months p=0.005, Student’s T-test). (I-K) Quantification of elongated soma of microglia in hippocampal CA2, CA3 and DG of 2- and 3-months old mice (n=3 each genotype, CA2 elongated p=0.027; CA3 elongated p=0.002; DG elongated p=0.018, Student’s T-test). (L-N) Quantification of CD68 foci per Ai14-labelled microglia in hippocampal CA2, CA3 and DG of 2- and 3-months old mice (n=3 each genotype, 2 months CA2 p=0.861, 3 months CA2 p=0.006; 2 months CA3 p=0.931, 3 months CA3 p=0.023; 2 months DG p=0.243, 3 months DG p=0.002, Student’s T-test). (O-P) Quantification of Iba1+ microglia in hippocampal CA2/3 and DG of 2- and 3-month-old control and ATRX miKO mice (n=3 each genotype, CA2/3 2 months p=0.975, 3 months p=0.00002; DG 2 months p=0.430, 3 months p=0.002, Student’s T-test). In all panels, control data is shown in black and ATRX miKO data is shown in red.

**Figure S2:**
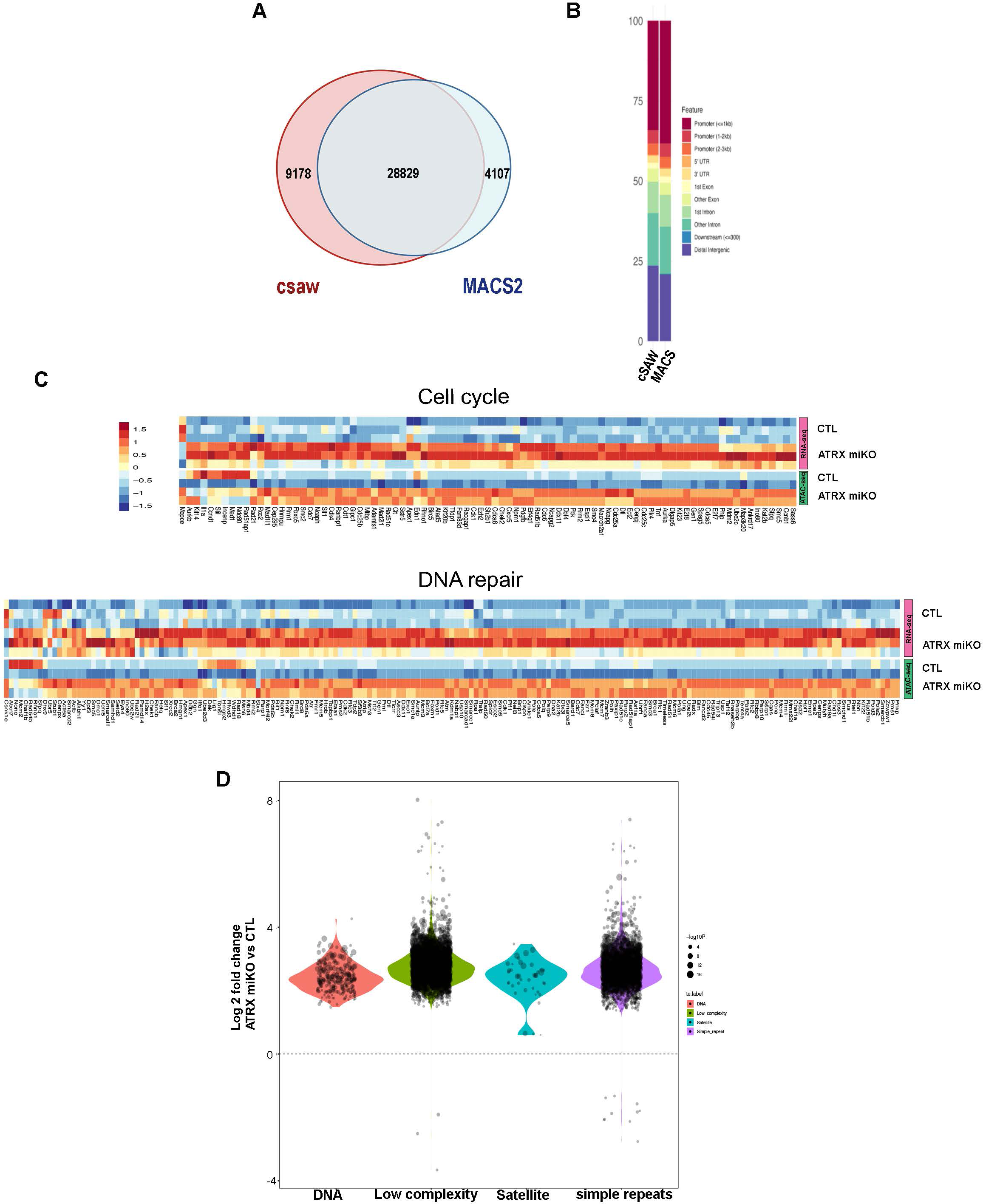
Chromatin accessibility in microglia lacking ATRX. (A) Overlap of DARs called by csaw and MACS. (B) Genomic distribution of DARs called by csaw and MACS. (C) Heatmaps represent an association between gene expression and chromatin accessibility for cell cycle and DNA repair pathways. The z-score was computed autonomously from the RNA expression or ATAC-seq signals. (D) Increased chromatin accessibility is observed at various types of repetitive elements in ATRX-null compared to control microglia.

**Figure S3:**
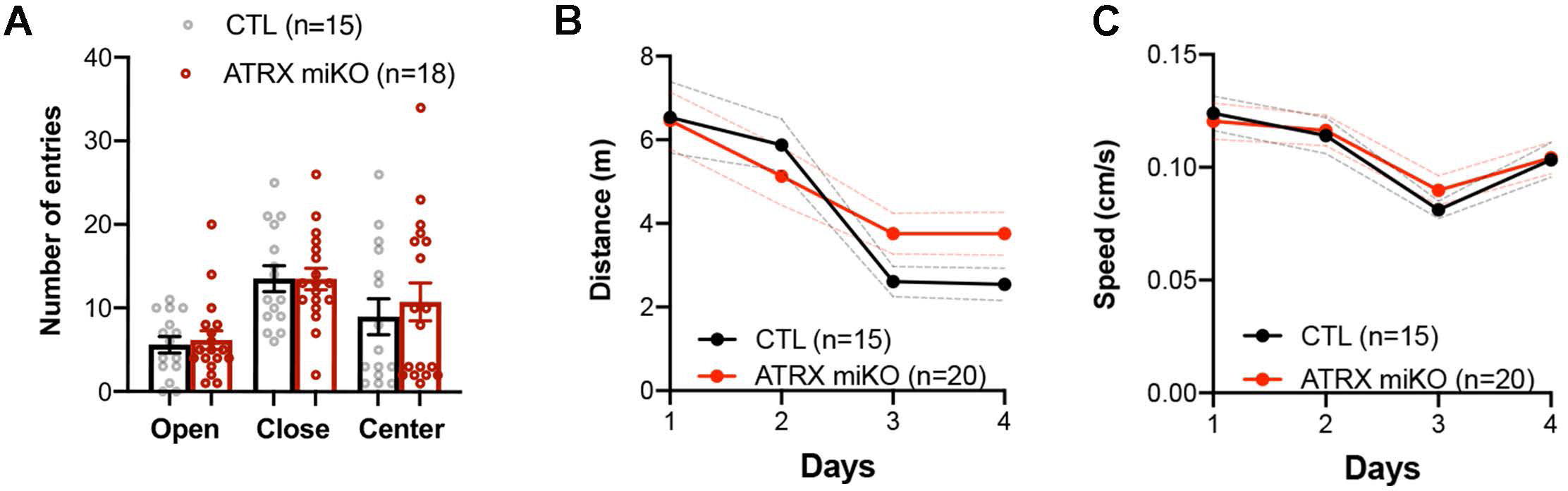
Supplemental data related to elevated plus maze and Morris water maze tests. (A) The number of entries in the open and closed arms or in the center of the elevated plus maze over 5 minutes (open arm entries p=0.710, closed arm entries p=0.986, center arm entries p=0.577, Student’s T-test). (B) Distance travelled and (C) swimming speed over 4 days of training (4 trails/day) in the Morris water maze task (distance travelled F(1, 33)=0.5162, p=0.477; swimming speed F(1, 132)=0.1696, p=0.681, two way ANOVA). Error bars represent ± SEM.

## Supplementary Tables

**Table S1:** Altered transcripts identified by RNA-seq of ATRX-null and control microglia nuclei

**Table S2:** Gene set enrichment analysis (GSEA) of altered transcripts

**Table S3:** DARs identified by ATAC-seq of ATRX-null and control microglia nuclei

**Table S4:** TOBIAS output from ATAC-seq data

**Table S5:** DARs at transposable elements (TEs)

**Table S6:** Altered transcripts at transposable elements (TEs)

**Table S7:** Cytokines/chemokines levels in *Atrx* miKO and control mice

**Table S8:** Oligonucleotides used in this study

